# T-Gene: Improved target gene prediction

**DOI:** 10.1101/803221

**Authors:** Timothy O’Connor, Charles E. Grant, Mikael Bodén, Timothy L. Bailey

## Abstract

**Motivation:** Identifying the genes regulated by a given transcription factor (its “target genes”) is a key step in developing a comprehensive understanding of gene regulation. Previously we developed a method for predicting the target genes of a transcription factor (TF) based solely on the correlation between a histone modification at the TF’s binding site and the expression of the gene across a set of tissues. That approach is limited to organisms for which extensive histone and expression data is available, and does not explicitly incorporate the genomic distance between the TF and the gene.

**Results:** We present the T-Gene algorithm, which overcomes these limitations. T-Gene can be used to predict which genes are most likely to be regulated by a TF, and which of the TF’s binding sites are most likely involved in regulating particular genes. T-Gene calculates a novel score that combines distance and histone/expression correlation, and we show that this score accurately predicts when a regulatory element bound by a TF is in contact with a gene’s promoter, achieving median positive predictive value (PPV) above 50%. T-Gene is easy to use via its web server or as a command-line tool, and can also make accurate predictions (median PPV above 40%) based on distance alone when extensive histone/expression data is not available for the organism. T-Gene provides an estimate of the statistical significance of each of its predictions.

**Availability:** The T-Gene web server, source code, histone/expression data and genome annotation files are provided at http://meme-suite.org.

## 1 Introduction

The regulation of the transcription of many genes involves the looping of distal chromatin regions bound by transcription factors (TFs) to bring them into contact with the gene’s promoter [7]. This contact activates or inhibits the action of transcriptional machinery at the transcription start site (TSS) the gene. These TF-bound chromatin regions function as “regulatory elements” (REs) in ways that are often unique to a specific cell type, condition, developmental stage or tissue (for brevity hereinafter referred to as a “tissue”). Defective binding of TFs to REs due to genomic mutations in the TF binding sites (e.g., “regulatory SNPs” [10], or in the TF itself [9] can cause dysregulation of genes and pathological phenotypes. The tissue-specific identification of REs and the TFs that bind to them is relatively easy compared to direct verification of the contacts they make with gene promoters. Thus, to decipher the genetic regulatory networks involved in most biological processes computational prediction methods are of significant value [17].

Although regulatory elements can be identified in several ways (see Yao *et al.* [17] for a review), in the current work we focus on REs bound by a particular transcription factor in a particular cell type/condition/tissue. The current preferred of determining where a TF binds the genome in a given tissue is to use the ChIP-seq assay—chromatin immunoprecipitation followed by sequencing [1]. This yields a set of genomic regions called “ChIP-seq peaks” that are highly-enriched for regions bound by the TF used in the assay. Our objective is primarily to predict the target genes of the TF under the conditions of the ChIP-seq experiment. However, since we will validate our predictions using Hi-C chromatin contact data [11], the method we describe should work equally with regulatory elements determined by other methods (such as the analysis of epigenetic signatures [18]), as long as the REs are similar in size to ChIP-seq peaks (typically about 100 bp).

The dominant approach for predicting the target genes of transcription factors is to simply use some function of the the genomic distance between the predicted transcription factor binding site (TFBS) and each gene promoter [14]. These methods are easy to use, and generally require only the locations of the ChIP-seq peaks, and a file containing the locations of the TSSes of the organism’s genes.

Underlying most distance-based methods is the assumption that each predicted TFBS regulates the closest gene, or that each gene is regulated by the closest TFBS, where distance is defined as the number of base-pairs between the TFBS and a TSS of the gene. However, a good deal of transcriptional regulation is via distal enhancer regions and involves chromatin looping [4] that bypasses the nearest gene. In one human cell line (GM12878), fully 41% of chromatin loops connecting a non-promoter region to a promoter skip one or more intervening promoters [11], violating the “closest gene” assumption. Similarly, if the target gene has multiple TSSs, distance-based methods cannot tell which TSS is the actual target of a TF bound at a nearby enhancer. Finally, if a TF binds at multiple locations near a gene, there is no guarantee that the closest binding site actually regulates the gene, as the “closest TFBS” method assumes.

We previously described a method—CisMapper [12]—for predicting the regulatory targets of TFs using TF ChIP-seq data and correlation between histone modifications and gene expression across a panel of tissues, and showed that it was more accurate than distance-based methods. That work built upon several prior methods for linking regulatory elements to target genes that are not based on distance alone. For example, the method of Ernst *et al.* [6] uses distance plus data for three histone modifications (H3K4me1, H3K4me2 and H3K27ac) and gene expression in a panel of tissues. It requires a supervised learning training step, and was not tested with regulatory elements predicted in a tissue not included in the panel. Similarly, Thurman *et al.* [16] showed that crosstissue correlation of DNaseI hypersensitivity (DHS) between DHS regions overlapping promoters DHS regions not overlapping promoters can predict regulatory relationships, but it is not clear how to extend their approach to linking TFBSes to promoters. DHS data is also available in far fewer organisms than histone modification data, restricting the applicability of that approach. The PreSTIGE algorithm [5] uses crosstissue correlation of H3K4me1 and expression, but it was designed for linking enhancers (not TFBSes) to genes, requires CTCF binding data, and only predicts regulatory links when both the H3K4me1 and expression signals are specifically enriched in a given tissue. He *et al.* [8] and Roy *et al.* [13] also proposed methods for training predictors of regulatory links between regulatory elements and genes using a large number of input features (e.g., histone modifications, DHS and TF ChIP-seq). These predictors are more accurate than the simple correlation-based approaches like PreSTIGE, but require data from many assays in order to make predictions in a tissue of interest. None of the prior methods (except CisMapper) was tested with TFBSes predicted by TF ChIP-seq.

The primary goal of the current work is to provide a method for analyzing peaks from TF ChIP-seq experiments that is as easy to use as existing distance-based methods, but is substantially more accurate. Our new computational method, T-Gene, combines distance and histone/expression correlation into a single score, and we show that it is substantially more accurate than CisMapper, makes more extensive predictions, produces calibrated statistical estimates, and can be used with or without expression and histone data, making it much more widely applicable. Researchers can use T-Gene via its web server or as a downloadable commandline tool, both of which are part of the MEME Suite (found at http://meme-suite.org). When used with TF ChIP-seq peak regions, T-Gene predictions can help answer two questions of significant interest to biologists studying gene regulation. Specifically, T-Gene’s predictions tell the user which genes are most likely to be regulated by the TF, and which of the TF’s binding sites are involved in regulating particular genes.

## 2 Methods

### 2.1 T-Gene command-line tool

The minimal input to the T-Gene command-line tool is a set of putative regulatory region (RE) coordinates (e.g., ChIP-seq peaks) for a given organism, and an annotation file specifying the TSSes of all known transcripts for the organism. The putative regulatory regions must be provided as a BED file, such as are output by ChIP-seq peak-calling programs (e.g., MACS [19]), and the gene annotation file must be in GTF format, such as those provided by Ensembl (http://ensembl.org/info/data/ftp/index.html).

As we shall show in the Results section, T-Gene can provide far more accurate predictions when provided with an (optional) tissue panel containing paired histone modification and gene expression data for a number of tissues in the organism. From the MEME Suite website, users can currently download three tissue panels (two human and one mouse) suitable for use with T-Gene, as well as annotation files for eight model organisms.

T-Gene constructs a putative regulatory “link” for all (RE, transcript) pairs whose genomic distance satisfies the maximum distance constraint, *D*, (*D* = 500,000 bp by default). T-Gene defines the distance between an RE and a transcript as the distance between the TSS of the transcript and the closest edge of the RE locus, or 0 if the RE locus overlaps the TSS. T-Gene labels each link as to whether it is a Closest-TSS (CT) or Closest-Locus (CL) link, or both. In the case where an RE or transcript would have no link, T-Gene adds a single link to the closest transcript or RE locus, respectively, assuring that every RE and every TSS has at least one link. T-Gene calculates up to five scores for each putative regulatory link. T-Gene always computes one score based solely on the distance between the RE locus and the TSS. T-Gene can also compute a more accurate score if the user provides auxiliary information in the form of a “tissue panel”—paired gene expression and histone modification data for a set of tissues in the organism. In that case, for each link, T-Gene also computes its histone/expression correlation, the statistical significance of the correlation, and, finally a statistical score that combines histone/expression correlation with RE-TSS distance. The computation of these five scores is described below.

For each link, T-Gene always computes the “Distance *p*-value”, which is designed to capture the tendency for regulatory links to be short—e.g., for transcription factors to bind near their gene targets. Consequently, the Distance *p*-value is defined as the probability that the link length (*d*) is as short or shorter than observed under a null model. For the null model, we assume that, given that an RE is observed within the maximum distance *D* of the TSS, it is equally likely to fall anywhere within that radius of the TSS. This suggests a uniform null model, so T-Gene defines the Distance *p*-value as *Pr*(*X* ≤ *d* |*X* ∼ *U* [0, *D*]). Since the RE can be either upstream or downstream of the TSS, T-Gene estimates the Distance *p*-value as (2*d* + *w*)*/*(2*D* + *w*), where *w* is the width of the RE in base-pairs. Note that this definition gives all links where the RE overlaps the TSS the same (minimal) Distance *p*-value of *w/*(2*D* + *w*), and the minimal Distance *p*-value decreases as the resolution of the RE data (e.g., ChIP-seq peak width) improves. Note also that links whose length exceeds *D* (e.g., some Closest-Locus or Closest-TSS links) are assigned a Distance *p*-value of 1.

The second score that T-Gene calculates for each link (when provided with a tissue panel) is the correlation between the level of a histone mark at the RE and the level of expression of the transcript across the tissues in the tissue panel. T-Gene log-transforms (*x′* = *log*(*x* + 1)) the histone and expression levels and then computes their Pearson correlation coefficient. In this work, we evaluate T-Gene using the histone mark H3K27ac, but the T-Gene website provides tissue panels for human and mouse that include both H3K27ac and H3K4me3 data. When provided with data for more than one histone mark in the tissue panel, T-Gene treats each (RE, transcript, histone) *triplet* as a separate link.

The third score computed by T-Gene is called the “Correlation *p*-value”, which estimates the statistical significance of the histone/expression correlation. To accurately estimate the *p*-value of the correlation, we propose a null model that breaks the relationship between the expression and histone values by randomly shuffling the order of the tissues for the expression data. T-Gene generates samples from this null distribution by shuffling the expression columns, computing the correlation for each link, and repeating the entire process ten times. (This generates ten random samples from the null correlation distribution for each link.) T-Gene then estimates the *p*-value of each observed correlation, using the complete set of null samples, as the fraction of null samples with correlations less than or equal to the observed correlation (with a pseudocount of 1 added to the numerator and the denominator of the fraction to avoid zeros).

The fourth T-Gene score combines the effects of histone/expression correlation and link length into a score that we refer to as the “CnD *p*-value” (short for Correlation and Distance *p*-value). This is score is simply the *p*-value of the *product* of the correlation and Distance *p*-values. This approach of combining evidence using the product of *p*-values has previously proven useful in different contexts in bioinformatics [2].

Finally, T-Gene converts either the CnD *p*-value or the Distance *p*-value into a *q*-value, depending on whether a tissue panel was provided or not, respectively. The *q*-value is defined as the minimum false discovery rate (FDR) required to consider this link statistically significant, using the method of Benjamini and Hochberg [3]. (See also Storey and Tibshirani [15] for genome-wide studies.) To keep the output format of T-Gene the same when used with or without a tissue panel, when T-Gene is used without a tissue panel it sets the correlation to 0, the Correlation *p*-value to 1, and the CnD *p*-value to the value of the Distance *p*-value for each link.

There are technical problems associated with transcripts that have very low expression levels across the tissue panel. For example, if both the expression and histone values are 0 for a link in all tissues but one, the resulting correlation score is 1, regardless of the actual expression level in that one tissue. If the maximum expression level is very low, it may be dominated by measurement noise, and empirical evidence suggests that such correlation scores are not good predictors of regulatory relationships. CisMapper deals with this problem by simply omitting such links, enforcing constraints on both the minimum expression value required in some tissue and the variation in expression across the tissue panel. Since we want T-Gene to be able to report links for all transcripts and REs, we take a different approach.

Firstly, T-Gene adds random Gaussian noise to all 0 values in both the histone and expression data in the tissue panel. The magnitude of this noise is a factor of ten smaller than the lowest non-zero value of the expression or histone level across the tissues. Note that if there is no non-zero expression of a TSS across the panel, the correlation score for the link will be 0 by definition.

Secondly, T-Gene can down-scale the computed correlation score of any link where the highest expression of the transcript in any tissue is less than *L*, a user-specified parameter that defaults to 0 (no scaling). Specifically, if the highest expression value for a given transcript is *E*, and *E < L*, the correlation scores of its links are all reduced by a factor of *E/L before* they are used in the computation of the Correlation *p*-values. Naturally, this same scaling is applied in computing the correlation of the null correlation scores as well in the Correlation *p*-value estimation process described above. We refer to *L* as the “Low Expression Correlation Adjustment Threshold” (LECAT).

The primary outputs of T-Gene are an interactive HTML report and a tab-separated values (TSV) file, each of which contains the links and their associated scores. The interactive HTML report allows the user to choose which fields to display (e.g., Gene ID, Gene Name, Strand, RE Locus, Distance, CT, CL, Correlation, CnD *p*-value etc.), to sort and filter the results based on any of those fields, and to download the displayed links as a TSV file. This allows the user to, for example, view (and save) the results sorted on the CnD *p*-value, or, alternatively, sorted on the Distance *p*-value. Using fields on the HTML output report, the user can also, for example, choose to view only links involving REs that are upstream (or downstream) of their putative target.

In addition, T-Gene’s HTML report and TSV file both include two fields (CT and CL) that specify if a particular link connects an RE to the closest TSS on its chromosome, or connects a transcript to its closest RE locus. As we show in the Results section, CT links prove to be more predictive of regulatory relationships than non-CT links. Consequently, by default the HTML report is sorted first on the CT field, followed by the CnD *p*-value (or Distance *p*-value if there is no tissue panel). This causes the CT links to appear before all other links, while ordering all links within the CT- and non-CT-link groups by statistical significance. The T-Gene output files also include fields giving the maximum expression value of the transcript and the maximum histone level of the RE across the tissue panel.

By default, the T-Gene HTML report and TSV file both include all CT (Closest-TSS) and CL (Closest-Locus) links, regardless of whether they violate the maximum distance constraint (*D*), and regardless of their *p*-values, as well as all links with CnD *p*-values no greater than 0.05. If the user desires that all links satisfying the maximum distance constraint be included, they can set the *p*-value threshold to 1.0 on the T-Gene command line. The user can also suppress CT- and CL-links that violate the distance constraint, if desired.

### 2.2 T-Gene web interface

The T-Gene web interface conveniently exposes most of the functionality of T-Gene. It allows the user to upload a file of regulatory elements in BED format and select a genome or a tissue panel from a drop-down list. Currently, the web interface provides two human tissue panels, one mouse tissue panel, and genomes for eight model organisms—arabidopsis (*A. thaliana*), worm (*C. elegans*), zebra fish (*D. rerio*) fly (*D. melanogaster*), human (*H. sapiens*), mouse (*M. musculus*), rat (*R. norvegicus*) and yeast (*S. cerevisiae*). The user can also adjust the maximum *p*-value for links to be reported and whether to include Closest-TSS and/or Closest-Locus links that violate the maximum distance constraint (*D*). The complete results of the T-Gene are returned as links to the HTML and TSV output files, as well as a compressed TAR file for more convenient downloading.

### 2.3 Testing methodology and data

#### 2.3.1 Evaluation

We validate the regulatory links predicted by T-Gene using a set of chromatin contacts predicted by the capture Hi-C (CHiC) assay in GM12878 [11]. This validation approach is based on the assumption that direct contact between a promoter and a distal chromatin region bound by a TF is required for the TF’s binding to affect expression of the gene. The CHiC assay captures Hi-C derived DNA fragments that contain a promoter region. The capture step enriches for chromatin contacts between promoter regions and other loci. We define the set of chromatin contacts *C* = {⟨*o, m*⟩}, where *o* and *m* are “other” and “promoter” regions from promoter-other CHiC contacts. We will refer to this as the “Mifsud” dataset.

Suppose that ⟨*p, t*⟩ is a predicted regulatory link, where *p* is a locus (e.g., a ChIP-seq peak) and *t* is a TSS. We say that this link is “confirmed” if it “coincides” with some contact ⟨*o, m*⟩ ∈ *C*. We say that link ⟨*p, t* ⟩coincides with contact ⟨*o, m*⟩ if locus *p* overlaps the “other” region *o*, and TSS *t* is contained in the promoter region *m*.

Using this rule, given a set of predicted regulatory links *L*, we determine the set of confirmed links *S*. As our estimate of prediction accuracy we compute the positive predictive value (PPV), which is the fraction of predicted links that are confirmed as true, 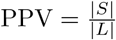. For plotting purposes we estimate the standard deviation of the PPV by assuming that the number of confirmed links |*S*| follows a binomial distribution. Thus, error bars are based on 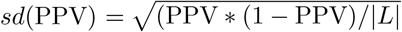. When plotting the average accuracy over a number of runs of T-Gene, we use boxplots of the PPV that show the range of the middle quartiles with a line at the median, and stars show outliers farther than 1.5 times the interquartile range (the whiskers) from the median.

The distribution of the distance between the *centers* of the two regions (*o* and *m*) for each contact in the Mifsud dataset is shown in Fig. 1. This data shows that the prior probability of a chromatin contact varies inversely with distance, suggesting that link length should be a good predictor of regulatory interactions.

**Figure 1:**
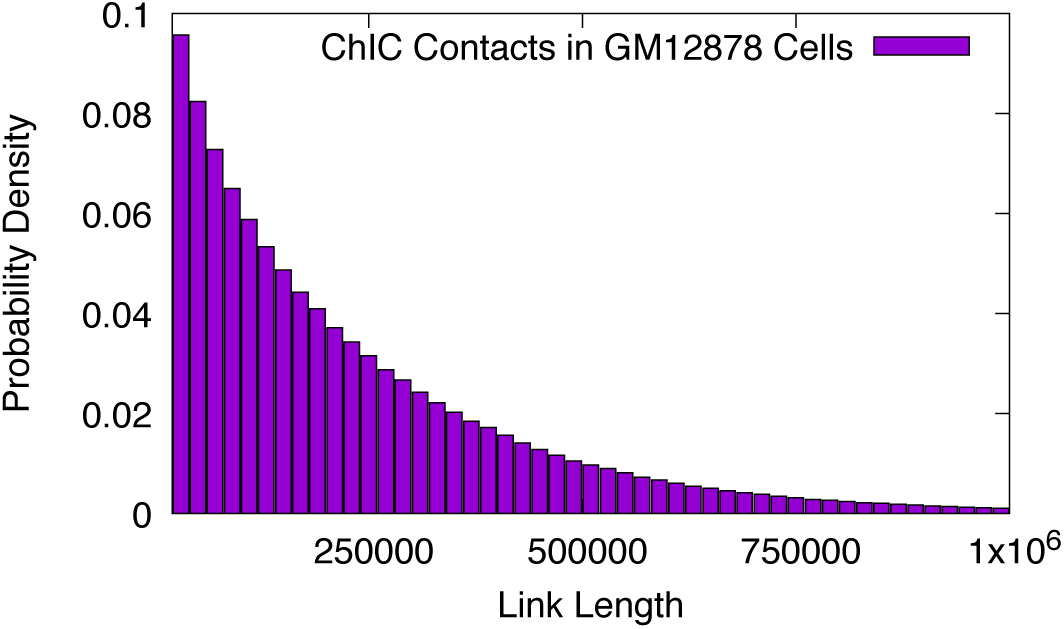
Length distribution of chromatin contacts in the Mifsud CHiC dataset. The plot shows the fraction of promoter-other CHiC contacts (y-axis) of a given length (x-axis) when considering only lengths up to 1 Mb in length.

Although not apparent from Fig. 1, the Mifsud dataset does not contain any contacts where the centers of the promoter and other regions are less than 20,000 bp apart (see Supplemental Fig. S1). For technical reasons explained in Mifsud *et al.* [11], they censored shorter contacts from their reported data. Consequently, we restrict our validation of T-Gene links to those whose length is at least 30,000 bp. This ensures that we do not attempt to validate regulatory links predicted by T-Gene where the (potential) validating contact was censored from the Mifsud data.

Unless otherwise noted, all experiments reported in the Results section use the default settings of the command-line version of T-Gene, with the exception of the value of LECAT, which is set to 6, as opposed to its default value of 0.

#### 2.3.2 Datasets

For evaluation studies, we downloaded 23 GM12878 transcription factor TF ChIP-seq datasets from the ENCODE repository at UCSC. We describe this in detail in Supplemental Table S1. Details on how we obtained the Mifsud *et al.* [11] CHiC contact data are given in the Supplement (in section “Data sources”). The data sources we used to construct our histone/expression tissue panels are described in Supplemental Table S2 (ENCODE 8-tissue panel) and Supplemental Table S3 (Roadmap Epigenomics 48-tissue panel).

## 3 Results

### 3.1 Accuracy of T-Gene regulatory link predictions

The relative merits of three of T-Gene’s scoring functions are illustrated in Fig. 2, which shows the results of running T-Gene using the Roadmap Epigenomics 48-tissue panel on 23 GM12878 TF ChIP-seq datasets. Averaged over the 23 sets of predictions made by T-Gene, the accuracy of the top links is substantially greater when links are sorted by CnD *p*-value, compared to sorting links by the correlation or Distance *p*-values (Fig. 2A). For example, the median PPV for the top 100 links is approximately 42% using CnD *p*-values, and slightly less than 30% using the other two scores. For comparison, randomly chosen links have a PPV of only about 15%. As shown in Fig. 2B, the CnD *p*-value performs best in the majority of cases, with the PPV of the 100 top links above 50% for several TFs. However, for six TFs (notably Pax5), sorting all links by Distance *p*-value gives higher PPV for the first 100 links.

**Figure 2:**
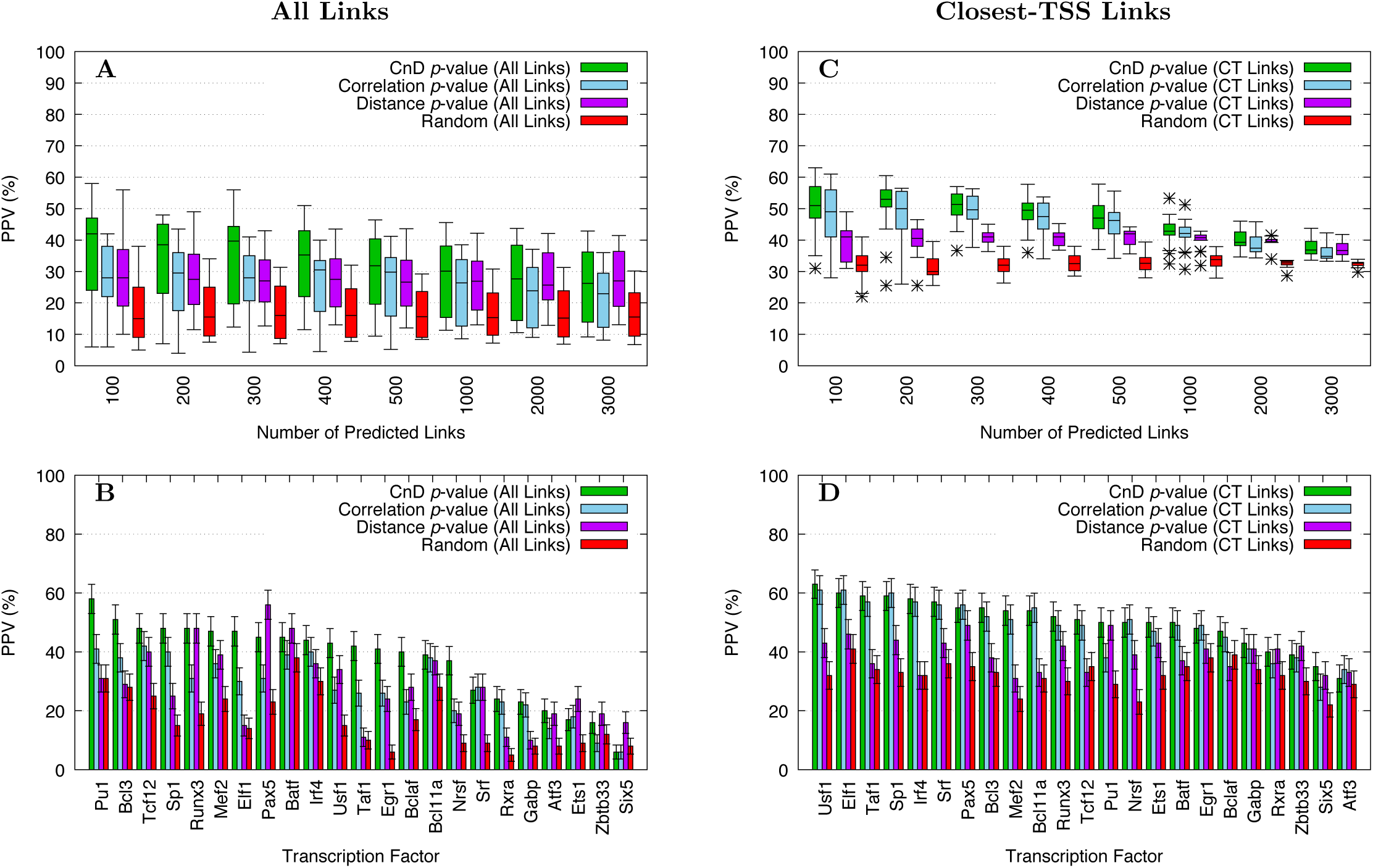
Accuracy of T-Gene scores using the Roadmap Epigenomics 48-tissue panel. Boxplots show the accuracy (PPV) of the top links (panel **A**) or top Closest-TSS links (panel **C**) predicted by T-Gene using the the CnD, Correlation or Distance *p*-values, averaged over 23 GM12878 TF ChIP-seq datasets. Barplots present the accuracy of T-Gene’s predictions on each of the 23 ChIP-seq datasets, focusing on the top 100 links (panel **B**), or the top 100 Closest-TSS links (panel **D**). The accuracy of randomly selected Closest-TSS links is shown for comparison (“Random”).

T-Gene achieves substantially higher predictive accuracy when we combine its Closest-TSS link filter with its scoring functions, as shown in Fig. 2C and Fig. 2D. The median PPV of the first 300 links is slightly more than 50% when the Closest-TSS links are sorted by CnD *p*-value (Fig. 2C). In this setting, the performance of the Correlation *p*-value is similar to that of the CnD *p*-value, reflecting the decreased additional information present in the link length when we restrict predictions to Closest-TSS links. Using the CnD *p*-value gives higher median PPV for (at least) the top 1000 predicted links, compared to using the Distance *p*-value, and similar performance up to the top 3000 links (Fig. 2C). As shown in Fig. 2D, with Closest-TSS links, the accuracy of the CnD *p*-value on the top 100 links is consistently as good or better than the other two T-Gene scores for all but one of the 23 TF ChIP-seq datasets, reaching PPV of approximately 60% for five TFs, and 50% or more for a majority of the TFs in this study.

Running T-Gene with the ENCODE 8-tissue panel rather than the Roadmap Epigenomics 48-tissue panel on the 23 GM12878 TF ChIP-seq datasets gives predicted links with very similar levels of accuracy (Fig. 3A). When T-Gene sorts all links using the CnD *p*-value, the median PPV of the top 100 links is 40%—approximately the same as when using the Roadmap Epigenomics tissue panel (compare with Fig. 2A). Accuracy is even better when T-Gene’s Closest-TSS filter is employed together with the CnD *p*-value, with PPV over 55% for the top 300 links (Fig. 3B). This increased accuracy may be due to the fact that the Roadmap Epigenomics expression data assigns the same expression value to all transcripts of a given gene, whereas the ENCODE data provides discrete expression values for each transcript. As with the Roadmap Epigenomics tissue panel, with the ENCODE 8-tissue panel T-Gene’s CnD *p*-value performs as well or better at prioritizing both all links and Closest-TSS links for up to 3000 predictions (Fig. 3A and B). The PPV of the top 100 Closest-TSS links sorted by CnD *p*-value is over 60% for five of the 23 TF ChIP-seq datasets (Tcf12, Elf1, Pax5, Bcl3 and Sp1), and it is over 50% for 14 of the datasets (Supplemental Fig. S2B).

**Figure 3:**
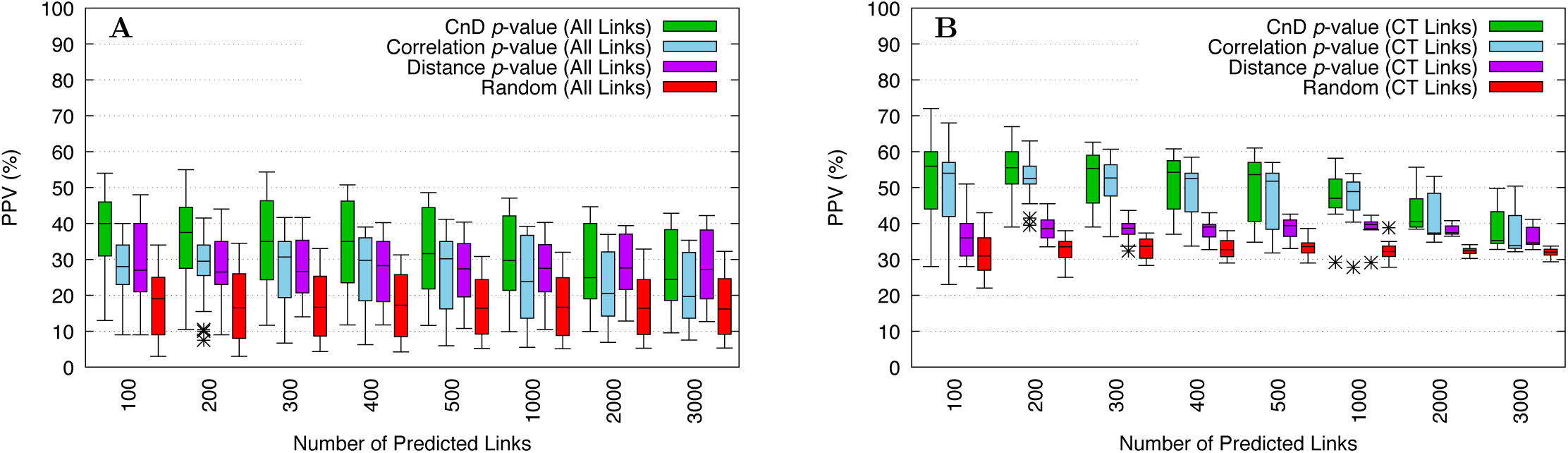
Accuracy of T-Gene scores using the ENCODE 8-tissue panel. Boxplots show the accuracy (PPV) of the top links (panel **A**) or top Closest-TSS links (panel **B**) predicted by T-Gene using the the CnD, Correlation or Distance *p*-values, averaged over 23 GM12878 TF ChIP-seq datasets. The accuracy of randomly selected Closest-TSS links is shown for comparison (“Random”).

We have shown that the subset of all links reported by T-Gene that are labeled by it as Closest-TSS links are considerably more reliable, especially when they are sorted according to the CnD *p*-value score. This is further illustrated in Fig. 4, where we compare the accuracy of links filtered by T-Gene on several criteria. Using T-Gene with either the Epigenetic Roadmap 48-tissue panel (Fig. 4A) or the ENCODE 8-tissue panel (Fig. 4B) leads to similar conclusions. Firstly, Closest-TSS (CT) links prove most accurate at all numbers of predicted links from 100 to 3000. Secondly, there is no synergy if we require that a link be both a Closest-Locus and a Closest-TSS link. Furthermore, the resulting subset of links is much smaller (hence the missing boxplots for “CL and CT Links” at the higher numbers of predicted links in Fig. 4). Thirdly, the overall accuracy of all links is dominated by that of the links that are not Closest-TSS links (non-CT links). Finally, Closest-Locus links are generally less accurate, on average, than the entire set of links. Thus, using T-Gene’s Closest-TSS filter seems extremely useful, but other filter combinations are not.

**Figure 4:**
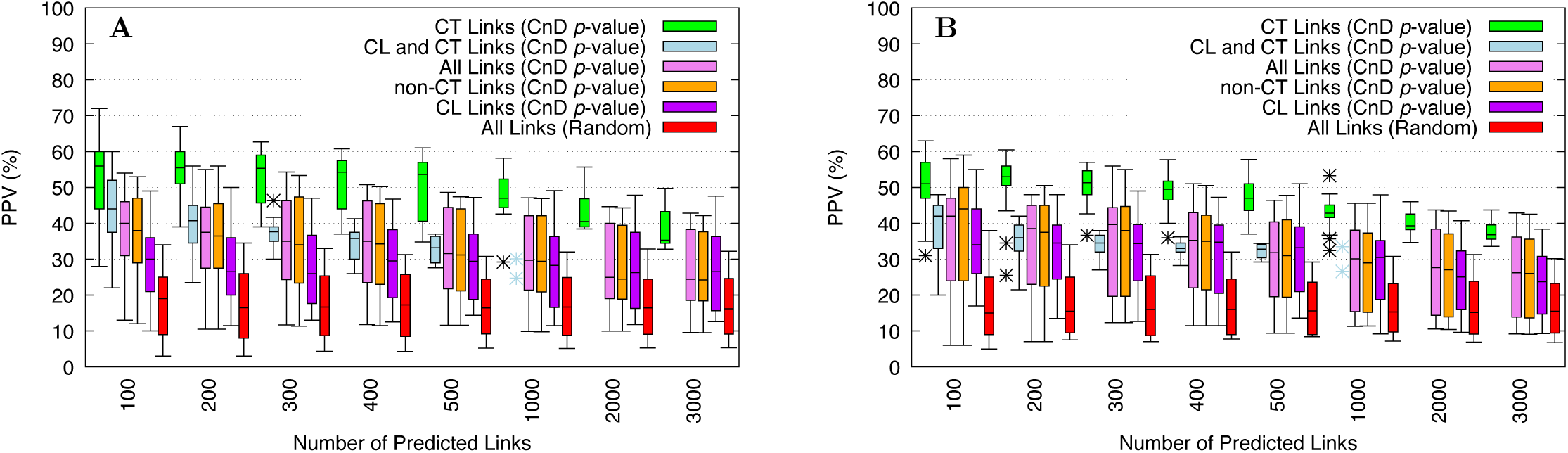
Accuracy of CnD *p*-value scores for different link types. Boxplots show the accuracy (PPV) of Closest-TSS links, all links, non-Closest-TSS links and Closest-Locus links, sorted by their CnD *p*-values, averaged over 23 GM12878 TF ChIP-seq datasets. T-Gene used the ENCODE 8-tissue panel (panel **A**), or the Roadmap Epigenomics 48-tissue panel (panel **B**), and LECAT=6. The accuracy of randomly selected links is shown for comparison (“Random”).

### 3.2 Reducing the effect of low gene expression

As we show in this section, when T-Gene computes the histone/expression correlation of a link, the value is less reliable if the expression of the transcript is uniformly low across the tissues in the tissue panel. Consequently, T-Gene allows the user to specify a value, *L*, called the “Low Expression Correlation Adjustment Threshold (LECAT)”, and T-Gene scales the computed correlation toward zero if the maximum expression of the transcript is less than *L*. To measure the effectiveness of this approach, we ran T-Gene using the ENCODE 8-tissue panel on the 23 GM12878 TF ChIP-seq datasets, while varying the value of *L* from 0 (no scaling) to 10 (scale correlation if maximum expression is less than 10).

Using a non-zero value of *L* can increase the median accuracy (PPV) of the first 100 links predicted by T-Gene from 30% (*L* = 0) to over 40% (*L* = 4), as shown in Fig. 5. The accuracy of the first *N* predicted links is higher using non-zero *L* even for the first 3000 links. The improvement is relatively insensitive to the size of *L* beyond a value of 4. When we repeat the same test using the Roadmap Epigenomics 48-tissue panel (Supplemental Fig. S4), using a non-zero value of *L* has very little effect. This may be due to the fact that there are fewer genes with very low expression values in the Roadmap Epigenomics data, which combines the expression at all TSSes of a gene into a single expression value for the gene. Based on these results, the T-Gene web server uses a value of *L* = 6 for the human and mouse tissue panels. The command-line version of T-Gene uses a default value of *L* = 0.

**Figure 5:**
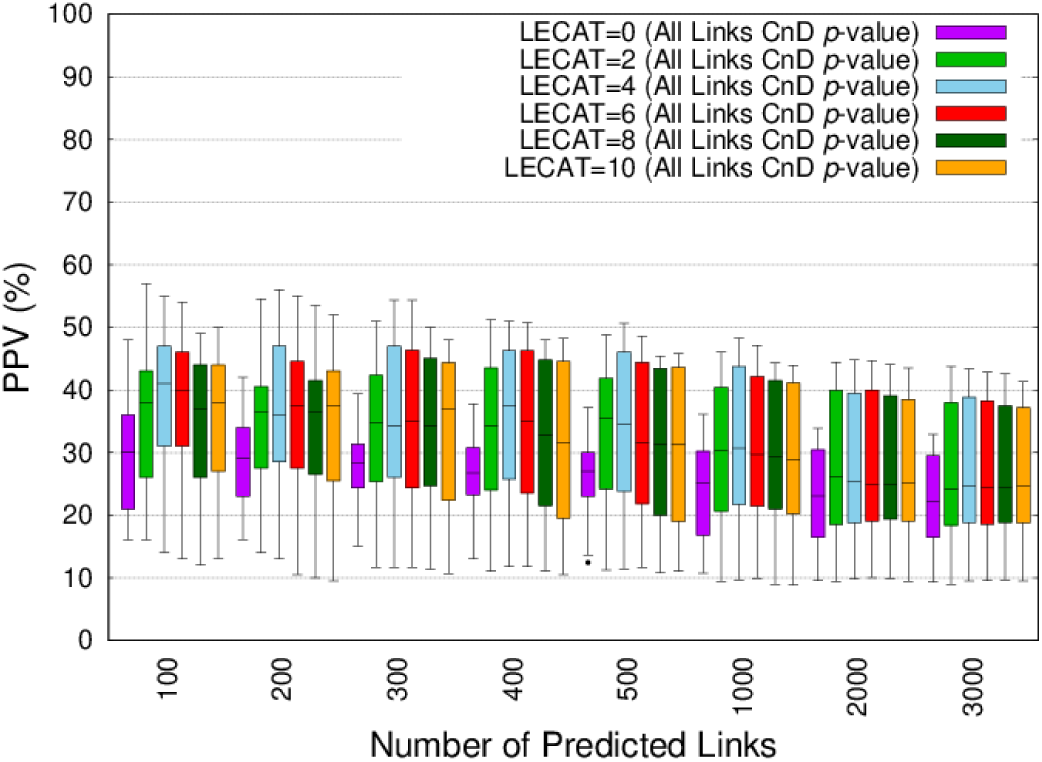
Effect of LECAT on the accuracy of T-Gene CnD *p*-values. Boxplots show the accuracy (PPV) of the top links predicted by T-Gene using the the CnD *p*-values with different values of the LECAT (Low Expression Correlation Adjustment Threshold) parameter, averaged over 23 GM12878 TF ChIP-seq datasets. T-Gene used the ENCODE panel of 8 tissues.

### 3.3 Statistical accuracy of T-Gene *p*-values

Fig. 6 shows Q-Q plots to assess the statistical accuracy of the *p*-values computed by T-Gene. We generated the plots by first creating a random set of ChIP-seq peaks from the largest of the 23 ENCODE GM12878 ChIP-seq datasets. For each of the 67,695 peaks in the original dataset, we created a random peak on the same chromosome with the same peak width, *w*. Next we created a random version of the ENCODE 8-tissue panel, where the order of the tissues is shuffled for the expression data, breaking any correspondence between expression and histone modification levels. We then ran T-Gene on the random peak file using the shuffled tissue panel. We then constructed a Q-Q plot by sorting the *p*-values reported by T-Gene for the *n* links, and for the *p*-value with rank *r* plotting (*X, Y*), where *Y* is the link’s *p*-value (as estimated by T-Gene), and *X* = *r/*(*n* + 1) is its “rank *p*-value”. If the reported *p*-values are statistically accurate, the *p*-values and rank *p*-values should be approximately equal for a given link, and the points will lie near the line *X* = *Y* in the Q-Q plot. Distance *p*-values are accurate except for very small values because, due to how we have defined them, they cannot be smaller than *w/*(2*D* + *w*) (Fig. 6A). This causes Distance *p*-values to be conservative (too large) for small distances. However, this is only noticeable for input files containing more than 1000 loci (Supplemental Fig. S3). Correlation *p*-values are accurate (Fig. 6B). Even with large numbers of loci (67,965 in this example) and despite the inaccuracy of the underlying Distance *p*-values for small distances, CnD *p*-values are only slightly conservative (Fig. 6C). As a result, T-Gene’s estimates of *q*-values are accurate under the assumptions of the null model we have defined (random peak positions and no relationship between expression and histone modification levels).

**Figure 6:**
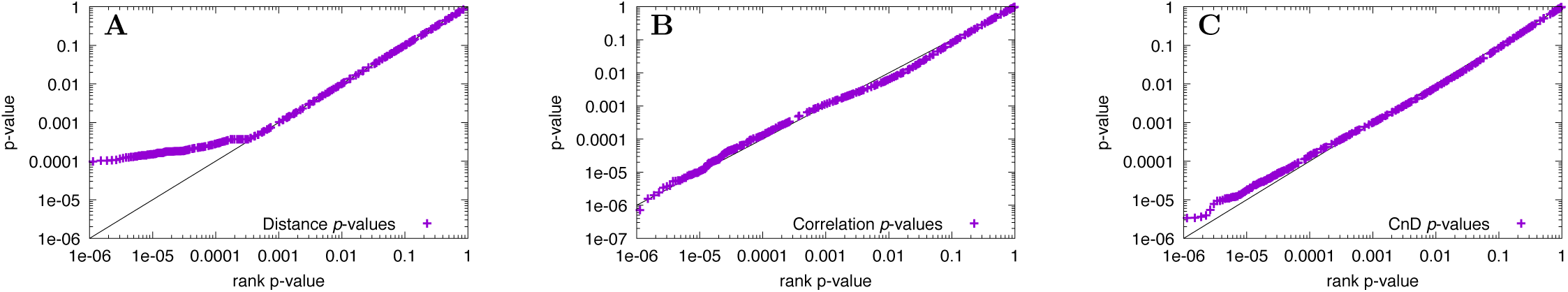
Statistical accuracy of T-Gene *p*-values. Q-Q plots show the accuracy of the *p*-values estimated by T-Gene on random input data for distance (panel **A**), expression-histone correlation (panel **B**) and the combination of correlation and distance (panel **C**). Each point (*X, Y*) represents one of *n* links, sorted by *p*-value, where *Y* is its *p*-value as estimated by T-Gene, and *X* = *r/*(*n* + 1), where *r* is the rank of its *p*-value. The diagonal line *X* = *Y* is shown for reference.

## 4 Discussion

We have presented a general-purpose algorithm for predicting regulatory relationships between genomic loci and genes. T-Gene can be used with sets of loci from any organism for which a gene annotation file exists. Such files can be downloaded for hundreds of species from ftp://ftp.ensembl.org/pub and ftp://ftp.ensemblgenomes.org/pub. T-Gene can be conveniently used via its web server at http://meme-suite.org/ `tgene`, or downloaded (along with gene annotation files and tissue panels), for use on the user’s computer. We currently provide tissue panels only for human and mouse, but we will continue to add to these as sufficient histone/expression data becomes available for other organisms. We provide gene annotation files for eight model organisms, and can easily add more in response to user requests.

Our results show that a combination of features is most effective for predicting chromatin contacts between regions bound by transcription factors and regions containing transcription start sites. Combining histone/expression correlation, the distance between the TF ChIP-seq peak and the TSS, and the fact that the TSS is the nearest-neighbor to the peak yields the most accurate predictions of chromatin contact, and, by inference, potential regulation of the gene by the TF. It is worth noting that the reciprocal situation—where the ChIP-seq peak is the closest one to the TSS is far less informative. This is perhaps not surprising, given that TFs tend to bind close to the genes that they regulate, whereas most genes are not regulated by a given TF.

Like our previous algorithm, CisMapper [12], T-Gene can use information from a tissue panel to compute a score based on the correlation between the histone modification state of the RE with the expression of the TSS. However, T-Gene improves on the CisMapper algorithm’s approach in five ways. First, T-Gene can be used to predict regulatory relationships in organisms where data for constructing a histone/expression panel is not available. This greatly increases T-Gene’s applicability. Second, since link length is known to correlate with the likelihood of a regulatory relationship, T-Gene computes a more predictive score that combines histone/expression correlation with link length. Third, unlike CisMapper, computes an accurate *p*-value for the correlation, which allows T-Gene to report the false discovery rate associated with each link via the *q*-value [3] statistic. A fourth improvement is that every TSS and every RE is included in at least one link in the T-Gene output, rather than omitting any TSS or RE whose shortest link would exceed the maximum link length constraint. Finally, T-Gene incorporates a heuristic that reduces false positives by increasing the influence of link length on the score of links where the transcript has very low expression across the tissue panel, rather than omitting such links entirely as CisMapper does.

## References

[1] Bailey, T., Krajewski, P., Ladunga, I., Lefebvre, C., Li, Q., Liu, T., Madrigal, P., Taslim, C., and Zhang, J. (2013). Practical Guidelines for the Comprehensive Analysis of ChIP-seq Data. PLoS Comput Biol, 9(11), e1003326.

[2] Bailey, T. L. and Gribskov, M. (1998). Combining evidence using p-values: application to sequence homology searches. Bioinformatics, 14(1), 48–54.

[3] Benjamini, Y. and Hochberg, Y. (1995). Controlling the False Discovery Rate: A Practical and Powerful Approach to Multiple Testing. Journal of the Royal Statistical Society. Series B (Methodological), 57(1), 289–300.

[4] Bulger, M. and Groudine, M. (2011). Functional and mechanistic diversity of distal transcription enhancers. Cell, 144(3), 327–339.

[5] Corradin, O., Saiakhova, A., Akhtar-Zaidi, B., Myeroff, L., Willis, J., Cowper-Sallari, R., Lupien, M., Markowitz, S., and Scacheri, P. C. (2014). Combinatorial effects of multiple enhancer variants in linkage disequilibrium dictate levels of gene expression to confer susceptibility to common traits. Genome Res, 24(1), 1–13.

[6] Ernst, J., Kheradpour, P., Mikkelsen, T. S., Shoresh, N., Ward, L. D., Epstein, C. B., Zhang, X., Wang, L., Issner, R., Coyne, M., Ku, M., Durham, T., Kellis, M., and Bernstein, B. E. (2011). Mapping and analysis of chromatin state dynamics in nine human cell types. Nature, 473(7345), 43–49.

[7] Farnham, P. J. (2009). Insights from genomic profiling of transcription factors. Nat Rev Genet, 10(9), 605–616.

[8] He, B., Chen, C., Teng, L., and Tan, K. (2014). Global view of enhancer-promoter interactome in human cells. Proc Natl Acad Sci U S A, 111(21), E2191–E2199.

[9] Ilsley, M. D., Gillinder, K. R., Magor, G. W., Huang, S., Bailey, T. L., Crossley, M., and Perkins, A. C. (2017). Krüppel-like factors compete for promoters and enhancers to fine-tune transcription. Nucleic acids research.

[10] Macintyre, G., Bailey, J., Haviv, I., and Kowalczyk, A. (2010). is-rSNP: a novel technique for in silico regulatory SNP detection. Bioinformatics, 26(18), i524–i530.

[11] Mifsud, B., Tavares-Cadete, F., Young, A. N., Sugar, R., Schoenfelder, S., Ferreira, L., Wingett, S. W., Andrews, S., Grey, W., Ewels, P. A., Herman, B., Happe, S., Higgs, A., LeProust, E., Follows, G. A., Fraser, P., Luscombe, N. M., and Osborne, C. S. (2015). Mapping long-range promoter contacts in human cells with high-resolution capture Hi-C. Nat Genet, 47(6), 598–606.

[12] O’Connor, T., Bodén, M., and Bailey, T. L. (2017). CisMapper: predicting regulatory interactions from transcription factor ChIP-seq data. Nucleic acids research, 45, e19.

[13] Roy, S., Siahpirani, A. F., Chasman, D., Knaack, S., Ay, F., Stewart, R., Wilson, M., and Sridharan, R. (2015). A predictive modeling approach for cell line-specific long-range regulatory interactions. Nucl Acids Res, 43(18), 8694–8712.

[14] Sikora-Wohlfeld, W., Ackermann, M., Christodoulou, E. G., Singaravelu, K., and Beyer, A. (2013). Assessing Computational Methods for Transcription Factor Target Gene Identification Based on ChIP-seq Data. PLoS Comput Biol, 9(11), e1003342.

[15] Storey, J. D. and Tibshirani, R. (2003). Statistical significance for genomewide studies. Proc Natl Acad Sci U S A, 100(16), 9440–9445.

[16] Thurman, R. E., Rynes, E., Humbert, R., Vierstra, J., Maurano, M. T., Haugen, E., Sheffield, N. C., Stergachis, A. B., Wang, H., Vernot, B., Garg, K., John, S., Sandstrom, R., Bates, D., Boatman, L., Canfield, T. K., Diegel, M., Dunn, D., Ebersol, A. K., Frum, T., Giste, E., Johnson, A. K., Johnson, E. M., Kutyavin, T., Lajoie, B., Lee, B.-K., Lee, K., London, D., Lotakis, D., Neph, S., Neri, F., Nguyen, E. D., Qu, H., Reynolds, A. P., Roach, V., Safi, A., Sanchez, M. E., Sanyal, A., Shafer, A., Simon, J. M., Song, L., Vong, S., Weaver, M., Yan, Y., Zhang, Z., Zhang, Z., Lenhard, B., Tewari, M., Dorschner, M. O., Hansen, R. S., Navas, P. A., Stamatoyannopoulos, G., Iyer, V. R., Lieb, J. D., Sunyaev, S. R., Akey, J. M., Sabo, P. J., Kaul, R., Furey, T. S., Dekker, J., Crawford, G. E., and Stamatoyannopoulos, J. A. (2012). The accessible chromatin landscape of the human genome. Nature, 489(7414), 75–82.

[17] Yao, L., Berman, B. P., and Farnham, P. J. (2015). Demystifying the secret mission of enhancers: linking distal regulatory elements to target genes. Critical reviews in biochemistry and molecular biology, 50, 550–573.

[18] Zentner, G. E., Tesar, P. J., and Scacheri, P. C. (2011). Epigenetic signatures distinguish multiple classes of enhancers with distinct cellular functions. Genome Res, 21(8), 1273–1283.

[19] Zhang, Y., Liu, T., Meyer, C. A., Eeckhoute, J., Johnson, D. S., Bernstein, B. E., Nussbaum, C., Myers, R. M., Brown, M., Li, W., and Liu, X. S. (2008). Model-based analysis of ChIP-Seq (MACS). Genome Biol, 9(9), R137.

