## Supplement for "T-Gene: Improved target gene prediction"

*To whom correspondence should be addressed.

#### 1 Supplemental Results

Supplemental Fig. S1 shows the (partial) distribution of the distance between the centers of the two regions ($o$ and $m$) for each contact in the Mifsud *et al.* [1] ChIC dataset for distances up to 1Mb. Only distances between 0 bp and 21000 bp are shown in the barplot. Note that there are no contacts with distances less than 20000 bp in the Mifsud data. This figure is based on the same data as Fig. 1 in the main paper.

Supplemental Fig. S2 shows barplots of the accuracy of the top 100 T-Gene links and CT-only links for each of the 23 GM12878 TF ChIP-seq datasets. This figure is based on the same data as the boxplots in Fig. 3 in the paper, which presents the *average* accuracy as a function of the number of predicted links.

Supplemental Fig. S3 shows the Q-Q plots for the Distance $p$-values and CnD $p$-values reported by T-Gene when run with sets of random ChIP-seq peaks derived from three actual ChIP-seq datasets of different sizes. The rightmost column presents the same data as Fig. 6A and Fig. 6C in the main paper.

Supplemental Fig. S4 shows accuracy of T-Gene CnD $p$-values with different settings of the "Low Expression Correlation Adjustment Threshold" (LECAT) parameter, which reduces the correlation score for target transcripts with uniformly low expression across the tissue panel. In this figure, T-Gene was run using the Roadmap 48-tissue panel, rather than using the ENCODE 8-tissue panel (Fig. 5 in the main paper).

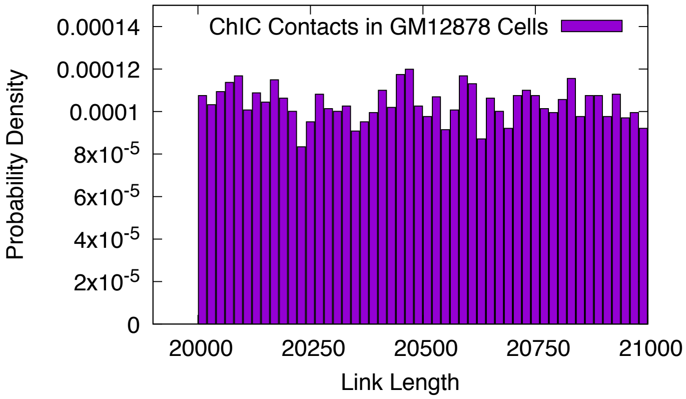


Figure S1: **Length distribution of chromatin contacts in the Mifsud CHiC dataset for links between 0 and 21,000 bp.** The plot shows the fraction of promoter-other CHiC contacts (y-axis) of a given length (x-axis) when considering only lengths up to 1 Mb in length.

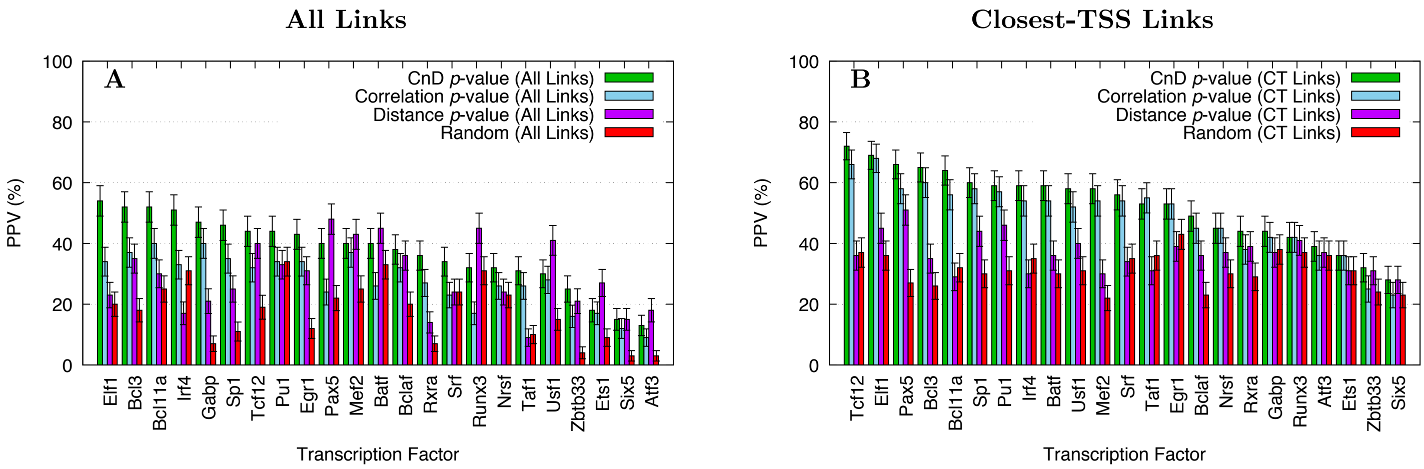


Figure S2: **Accuracy of T-Gene scores using the ENCODE 8-tissue panel.** Barplots present the accuracy of T-Gene's predictions on each of the 23 ChIP-seq datasets, focusing on the top 100 links (panel **A**), or the top 100 Closest-TSS links (panel **B**). The accuracy of randomly selected Closest-TSS links is shown for comparison ("Random").

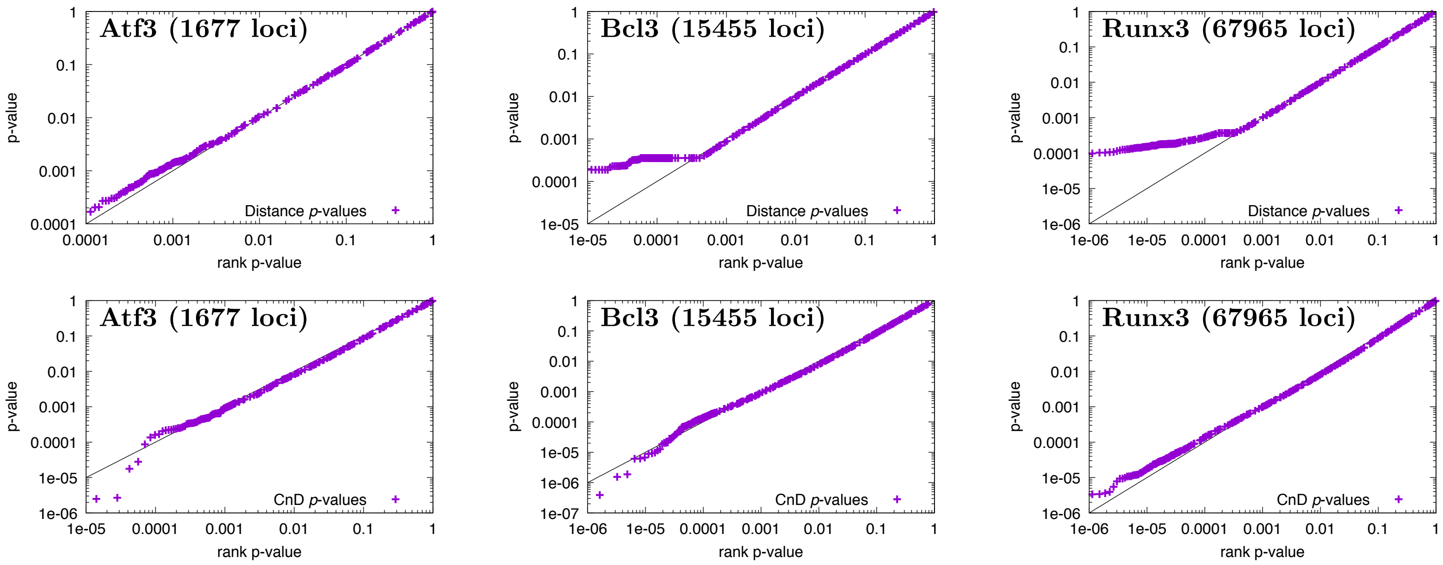


Figure S3: **Statistical accuracy of T-Gene $p$-values as a function of input size.** Q-Q plots show the accuracy of the Distance $p$-values (row 1) and CnD $p$-values (row 2) reported by T-Gene on sets of different numbers of random loci (see main paper for a description of the randomization method). Each point $(X, Y)$ represents one of $n$ links, sorted by $p$-value, where $Y$ is its $p$-value as estimated by T-Gene, and $X = r/(n + 1)$, where $r$ is the rank of its $p$-value. The diagonal line $X = Y$ is shown for reference.

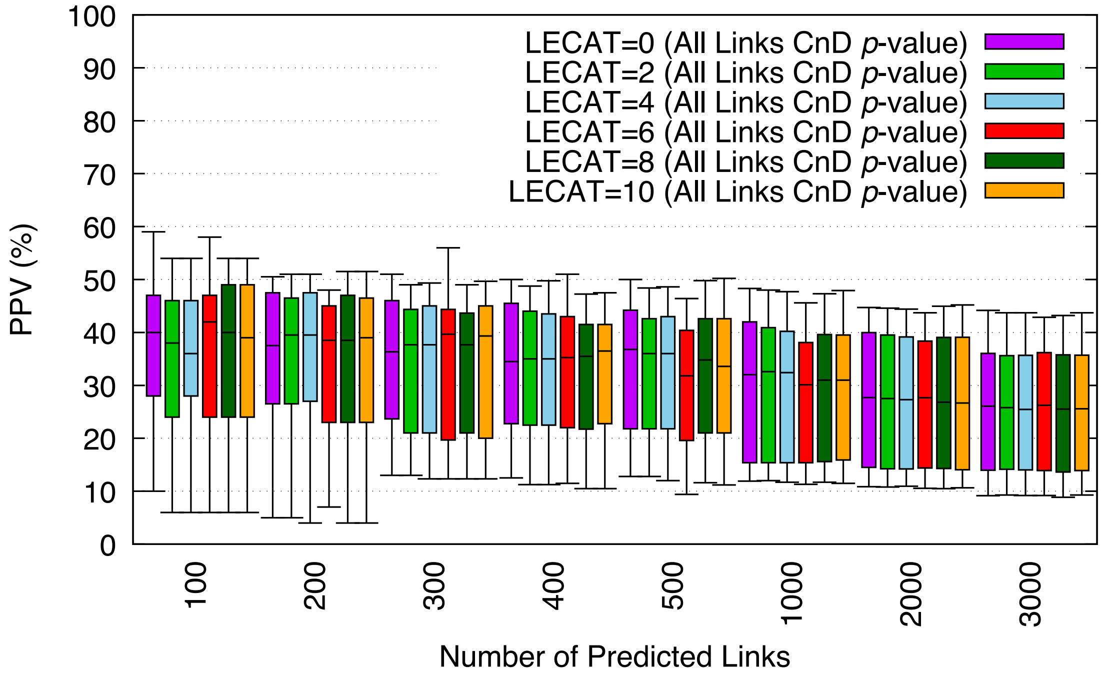


Figure S4: **Effect of LECAT on the accuracy of T-Gene CnD *p*-values (Roadmap 48-tissue panel).** Boxplots show the accuracy (PPV) of the top links predicted by T-Gene using the the CnD *p*-values with different values of the LECAT (Low Expression Correlation Adjustment Threshold) parameter, averaged over 23 GM12878 TF ChIP-seq datasets. T-Gene used the Roadmap Epigenomics panel of 48 tissues.

| *TF* | *File Name* | Number of Peaks | *TF* | *File Name* | Number of Peaks |
|---|---|---|---|---|---|
| Atf3 | Atf3Pcr1x | 1677 | Pax5 | Pax5c20Pcr1x | 25342 |
| Batf | BatfPcr1x | 32427 | Pu1 | Pu1Pcr1x | 42938 |
| Bcl11a | Bcl11aPcr1x | 17876 | Runx3 | Runx3sc101553V0422111 | 67965 |
| Bcl3 | Bcl3V0416101 | 15455 | Rxra | RxraPcr1x | 1704 |
| Bclaf | Bclaf101388V0416101 | 6114 | Six5 | Six5Pcr1x | 4839 |
| Egr1 | Egr1Pcr2x | 16331 | Sp1 | Sp1Pcr1x | 18248 |
| Elf1 | Elf1sc631V0416101 | 23008 | Srf | SrfPcr2x | 8544 |
| Ets1 | Ets1Pcr1x | 4120 | Taf1 | Taf1Pcr1x | 14278 |
| Gabp | GabpPcr2x | 6566 | Tcf12 | Tcf12Pcr1x | 20437 |
| Irf4 | Irf4sc6059Pcr1x | 17771 | Usf1 | Usf1Pcr2x | 9778 |
| Mef2a | Mef2aPcr1x | 17605 | Zbtb33 | Zbtb33Pcr1x | 2144 |
| Nrsf | NrsfPcr1x | 6906 | | | |

Table S1: **ENCODE TF ChIP-seq datasets.** The transcription factor ChIP-seq data used in this work was all downloaded from `http://hgdownload.soe.ucsc.edu/goldenPath/hg19/encodeDCC/wgEncodeAwgTfbsUniform`. The ChIP-seq peak file for each TF named in the first column of the table was downloaded by appending the following filename to the URL, where [File] is replaced by the name in the “File Name” column of the table: `wgEncodeAwgTfbsHaibGm12878[File]UniPk.narrowPeak.gz`.

| ENCODE Expression Data Sources | |
|---|---|
| Directory Name: | wgEncodeRikenCage/ |
| Filename format: | wgEncodeRikenCage[CellLine]CellPapTssGencV7.gtf.gz |
| Cell Lines: | Ag04450, Gm12878, H1hesc, Helas3, Hepg2, Huvec, K562, Nhek |

| ENCODE Histone Modification Data Sources | |
|---|---|
| Directory Name: | wgEncodeBroadHistone/ |
| Filename format: | wgEncodeBroadHistone[CellLine][Modification]StdPk.broadPeak.gz |
| Histone Mark: | H3k27ac |
| Cell Lines: | Ag04450, Gm12878, H1hesc, Helas3, Hepg2, Huvec, K562, Nhek |

Table S2: **ENCODE tissue panel data.** All histone and expression data for the ENCODE 8-tissue panel was downloaded from files in the named directory under `http://hgdownload.soe.ucsc.edu/goldenPath/hg19/encodeDCC/`, with [CellLine] replaced in the "Filename format" by one of the "Cell Lines" in the table above.

| Tissue ID | Tissue Name | Tissue ID | Tissue Name |
|---|---|---|---|
| E003 | H1_Cell_Line | E085 | Fetal_Intestine_Small |
| E004 | H1_BMP4_Derived_Mesendoderm_Cultured_Cells | E087 | Pancreatic_Islets |
| E005 | H1_BMP4_Derived_Trophoblast_Cultured_Cells | E094 | Gastric |
| E006 | H1_Derived_Mesenchymal_Stem_Cells | E095 | Left_Ventricle |
| E007 | H1_Derived_Neuronal_Progenitor_Cultured_Cells | E096 | Lung |
| E011 | hESC_Derived_CD184+_Endoderm_Cultured_Cells | E097 | Ovary |
| E012 | hESC_Derived_CD56+_Ectoderm_Cultured_Cells | E098 | Pancreas |
| E013 | hESC_Derived_CD56+_Mesoderm_Cultured_Cells | E100 | Psoas_Muscle |
| E016 | HUES64_Cell_Line | E104 | Right_Atrium |
| E037 | CD4_Memory_Primary_Cells | E105 | Right_Ventricle |
| E038 | CD4_Naive_Primary_Cells | E106 | Sigmoid_Colon |
| E047 | CD8_Naive_Primary_Cells | E109 | Small_Intestine |
| E050 | Mobilized_CD34_Primary_Cells_Female | E112 | Thymus |
| E055 | Penis_Foreskin_Fibroblast_Primary_Cells_skin01 | E113 | Spleen |
| E056 | Penis_Foreskin_Fibroblast_Primary_Cells_skin02 | E114 | A549 |
| E058 | Penis_Foreskin_Keratinocyte_Primary_Cells_skin03 | E116 | GM12878 |
| E059 | Penis_Foreskin_Melanocyte_Primary_Cells_skin01 | E117 | HELA |
| E061 | Penis_Foreskin_Melanocyte_Primary_Cells_skin03 | E118 | HEPG2 |
| E062 | Peripheral_Blood_Mononuclear_Primary_Cells | E119 | HMEC |
| E065 | Aorta | E120 | HSMM |
| E066 | Adult_Liver | E122 | HUVEC |
| E071 | Brain_Hippocampus_Middle | E123 | K562 |
| E079 | Esophagus | E127 | NHEK |
| E084 | Fetal_Intestine_Large | E128 | NHLF |

Table S3: **Roadmap Epigenomics tissue panel data.** All the data in the Roadmap Epigenomics 48-tissue panel was downloaded from `http://egg2.wustl.edu`. The first column in the table shows the Roadmap Epigenomics Project identifier (ID) for the tissue or cell line named in the second column. The expression data for the tissue is contained in a column with the given tissue ID in the expression data file that was downloaded from URL: `http://egg2.wustl.edu/roadmap/data/byDataType/rna/expression/57epigenomes.RPKM.pc.gz`. The H3K27ac ChIP-seq file for a given tissue was obtained by replacing [ID] in the following URL with the tissue's ID: `http://egg2.wustl.edu/roadmap/data/byFileType/peaks/consolidated/broadPeak/[ID]-H3K27ac.broadPeak.gz`.

#### 2 Data sources

From URL `http://www.ebi.ac.uk/arrayexpress/files/E-MTAB-2323/E-MTAB-2323.additional.1.zip` we obtained the Mifsud *et al.* [1] GM12878 CHiC chromatin contact data file: `TS5_GM12878_promoter-other_significant_interactions.tsv`, which is contained in the ZIP file.

Supplemental Table S1 describes how we downloaded the 23 GM12878 transcription factor ChIP-seq datasets that we used in evaluating the accuracy of T-Gene links.

The data sources for the histone/expression tissue panels used in this work are detailed in Supplemental Table S2 (8-tissue ENCODE panel) and Supplemental Table S3 (48-tissue Roadmap Epigenomics panel).

### References

[1] Mifsud, B., Tavares-Cadete, F., Young, A. N., Sugar, R., Schoenfelder, S., Ferreira, L., Wingett, S. W., Andrews, S., Grey, W., Ewels, P. A., Herman, B., Happe, S., Higgs, A., LeProust, E., Follows, G. A., Fraser, P., Luscombe, N. M., and Osborne, C. S. (2015). Mapping long-range promoter contacts in human cells with high-resolution capture Hi-C. *Nat Genet*, **47**(6), 598–606.
